# Comprehensive evaluation of the antimicrobial properties of platelet-rich fibrin *in vitro* in the context of the oral microbiome and bacterial species diversity

**DOI:** 10.1101/2025.05.28.656546

**Authors:** Wojciech Popowski, Dominika Domanowska, Damian Koseski, Rafał Ostrowski, Magdalena Zalewska, Milena Małecka-Giełdowska, Anna Łasica, Magdalena Popowska

## Abstract

Platelet-rich fibrin (PRF) is a platelet concentrate widely applied in various medical fields and is considered a valuable adjunct in tissue regeneration during surgical procedures. However, infections caused by biofilm-forming bacteria at surgical sites, combined with increasing antibiotic resistance, present a major clinical concern. Current research is focused on identifying alternative therapeutic strategies to improve infection control and promote wound healing. This study aimed to characterize the oral microbiome of healthy individuals and evaluate the *in vitro* antimicrobial properties of two PRF formulations. The antibacterial activity, along with its temporal dynamics at different initial bacterial concentrations, was assessed against Gram-negative bacteria (*Escherichia coli*, *Porphyromonas gingivalis*) and Gram-positive bacteria exhibiting diverse morphologies (*Bacillus subtilis*, *Micrococcus luteus*, *Staphylococcus lentus*, *Enterococcus casseliflavus*, *Streptococcus mutans*). Our results fill gaps in knowledge concerning the spectrum of PRF’s antimicrobial activity, demonstrating efficacy against a range of opportunistic and pathogenic bacteria. Key findings include the absence of significant differences in oral microbiome composition between male and female participants, a lack of inhibitory effect of A-PRF against *S. mutans*, and a transient inhibitory effect against *P. gingivalis* observed only at low initial OD₆₀₀ and within 24 hours. These results suggest that A-PRF therapy should be considered only in patients without active oral infection.

## 1. Introduction

Platelet concentrates are blood-derived products obtained by fractionating plasma through centrifugation using specialized equipment. By adjusting centrifugation parameters, various types of platelet concentrates can be produced, which are classified based on their cellular composition and the architecture of the fibrin matrix^1,2^.

The development of platelet concentrate preparations dates back to the 1970s, when Matras conducted pioneering research on fibrin glues to enhance wound healing in rat skin. In the following years, several studies proposed improved methods for utilizing blood-derived preparations, achieving a higher concentration of platelets in the final product. These techniques represented the first generation of platelet-rich plasma (PRP) gels. The application of these preparations yielded promising outcomes in fields such as ophthalmology, neurosurgery, and general surgery^3^. Following the studies by Whitman in 1997, previously developed preparations began to be widely used in oral and maxillofacial surgery as well as in regenerative medicine. These products were again collectively referred to as PRP, but without precise characterization of their cellular content or fibrin architecture^3^. During the same period, an alternative form of platelet concentrate, termed platelet-rich fibrin (PRF), was developed in France. The method of PRF preparation differed substantially from earlier techniques and was therefore classified as a second-generation platelet concentrate^3,4^. The first generation of platelet concentrates includes PRP with low leukocyte content, which was burdened with impaired wound healing due to the presence of anticoagulants and a detrimental effect on the cellular inflammatory response^1^. The second generation— PRF — is free of anticoagulants, thereby eliminating the adverse effects observed with first-generation platelet concentrates. Variants within this category include leukocyte- and platelet-rich fibrin (L-PRF), advanced platelet-rich fibrin (A-PRF), and injectable platelet-rich fibrin (i-PRF)^1^. Each type of PRF requires specific centrifugation parameters. Standard PRF is centrifuged at 700×g (approximately 4550 rpm) for 12 minutes, whereas advanced A-PRF is prepared at 200×g (approximately 1300 rpm) for 14 minutes, and i-PRF is obtained by centrifugation at 60×g (approximately 390 rpm) for 3 minutes^1,5,6^.

PRF contains a variety of biologically active molecules, including insulin-like growth factor 1 (IGF-1), platelet-derived growth factor (PDGF), vascular endothelial growth factor (VEGF), fibroblast growth factor (FGF), epidermal growth factor (EGF), platelet-derived epidermal growth factor (PDEGF), transforming growth factor beta (TGF-β), as well as proteins of the fibrin matrix, all of which are present at higher concentrations than in peripheral blood^1,7^. Among these, PDGF, VEGF, and TGF-β play essential roles in angiogenesis and the formation of new tissue structures^7,8^. The platelets within PRF contain granules loaded with cytokines, chemokines, and other inflammatory mediators that, upon release, enhance hemostasis and promote the activation and recruitment of cells to sites of inflammation. Furthermore, leukocytes present in PRF contribute to both angiogenesis and lymphangiogenesis through intercellular interactions and the expression of various signaling molecules^7^. Moreover, PRF exhibits osteoinductive, anti-inflammatory, pro-angiogenic, and antibacterial properties^9–11^. Platelet concentrates are widely utilized to induce and stimulate tissue repair and regeneration processes^1^.

In recent years, autologous platelet concentrates have been widely used in dentistry, maxillofacial surgery, and plastic surgery^9,12^. PRF has been applied in various procedures, including maxillary sinus augmentation, bone defect regeneration around implants, guided bone regeneration (GBR), healing of post-extraction sockets, healing of bone defects following enucleation of extensive periapical cysts, temporomandibular joint disorder treatment, healing of wounds in patients after oral cancer resection surgery, and periodontal surgery^4,10,13–19^. The use of PRF in sinus-lift procedures prior to planned implant placement can reduce healing time to 4 months; however, larger-scale studies are still needed to confirm this effect^4^. Current evidence supporting the necessity of adding PRF to bone grafts in sinus-lift procedures is limited^10^. PRF appears to be more effective in alleviating pain in patients with temporomandibular joint disorders, but additional research is required to fully determine the efficacy of such treatment^16^. The application of PRF in patients undergoing oral cancer resection surgery seems promising. In these patients, PRF improves tissue regeneration, reduces postoperative discomfort, and enhances treatment outcomes. Despite encouraging findings, further high-quality, randomized, controlled clinical trials are needed^17^. PRF has demonstrated a beneficial effect in reducing pain, swelling, and the incidence of osteitis following the extraction of impacted lower third molars^20^.

An increasing incidence of postoperative staphylococcal infections has been observed, leading to prolonged hospitalizations^9,21^. I-PRF, due to its content of proteins with antibacterial properties, may contribute to reducing the risk of such infections^9^. *In vitro* studies available in the literature indicate that both L-PRF and high-density PRF (H-PRF) exert antibacterial activity against *Staphylococcus aureus* and *Escherichia coli* strains^11^.

The another group of bacteria which may contribute to numerous diseases including post-treatment complications is oral microbiome. The human oral cavity comprises diverse habitats—including the inner cheeks, palate, tongue, and teeth—each supporting distinct microbial communities^22^. The oral microbiome is the second most diverse microbial community after the gut and includes bacteria, viruses, fungi, protozoa, and archaea^22,23^. Among these, bacteria are the most extensively studied and are found both in saliva and on oral surfaces such as mucosa, tongue, and teeth^22^. At the phylum level, a healthy oral microbiome is dominated by *Actinobacteria, Fusobacteria, Proteobacteria, Firmicutes*, and *Bacteroidetes*, which together account for approximately 96% of all oral bacteria^24,25^. *Fusobacteria* and *Bacteroidetes* are commonly cultivable, with *Fusobacteria* being among the most prevalent^22,25^. On the genus level, the microbiome is relatively stable; 11 genera are shared by over 99% of individuals and collectively make up about 77.8% of the total microbial abundance. These dominant genera include *Streptococcus, Prevotella, Veillonella, Lactobacillus, Actinomyces,* and *Neisseria*^22^. While most individuals share similar genera, species- and strain-level diversity remains highly individual-specific^25^. The core oral microbiome includes *Actinomyces, Atopobium, Corynebacterium,* and *Rothia* (*Actinobacteria*); *Bergeyella, Capnocytophaga*, and *Prevotella* (*Bacteroidetes*); *Granulicatella, Streptococcus*, and *Veillonella* (*Firmicutes*); *Campylobacter, Cardiobacterium, Haemophilus*, and *Neisseria* (*Proteobacteria*); as well as members of TM7 and *Fusobacteria*. Among *Firmicutes, Streptococcus* (19.2%) and *Veillonella* (8.6%) are the most abundant genera^24,26^. In contrast, the variable microbiome comprises genera that are less consistently present, such as *Chryseobacterium, Anaeroglobus, Filifactor, Lactobacillus, Johnsonella, Shuttleworthia*, unclassified OD2, *Brachymonas, Propiniovibrio, Scardovia, Olsenella*, and *Cryptobacterium*^24,26^. Due to constant exposure to external factors, the oral microbiota is subjected to dynamic fluctuations. Bacteria exhibit clear niche specificity: *Firmicutes* and *Actinobacteria* are predominantly found in dental plaque^27^, whereas anaerobes such as *Prevotella, Capnocytophaga*, and *Flavobacterium* (*Bacteroidetes*) colonize the dorsal and lateral surfaces of the tongue and microaerophilic niches^24^.

Fungi also constitute a notable component of the oral microbiome. Over 100 fungal species have been identified in healthy individuals^28^, with *Candida* spp. being the most prevalent, frequently contributing to early stages of biofilm formation^29^.

The diversity and stability of the oral microbiome are essential for oral and systemic health. Dysbiosis has been linked to conditions such as caries, periodontitis, and systemic diseases^22,25^. Periodontal disease arises from pathogenic biofilms that trigger gingival inflammation. Notably, certain members of the core microbiome, albeit in low abundance in healthy people,—*Porphyromonas gingivalis, Tannerella forsythia*, and *Treponema denticola*—form the so-called “red complex” and are strongly associated with periodontitis^24,30,31^. Risk factors that exacerbate disease progression include lack of proper hygiene, smoking, diabetes, obesity, and osteoporosis. Colonization of the periodontal pocket by pathogenic microorganisms promotes inflammation and tissue destruction^31^.

Although cultivation-based methods have identified many bacterial taxa, a large portion of oral microbes remains unculturable^23^. The expanded Human Oral Microbiome Database (eHOMD) offers curated information on bacterial taxa inhabiting the human oral cavity and aerodigestive tract. Among the 834 listed taxa, 523 are primarily oral and 22 are primarily nasal. Of the oral taxa, 49% are formally named, 21% are cultivated but unnamed, and 29% are known solely as uncultivated phylotypes^32^. To study both culturable and unculturable members of the microbiota, nucleic acid-based techniques such as 16S rRNA gene sequencing and shotgun metagenomics are commonly used. The former targets specific hypervariable regions for taxonomic resolution, while the latter offers a broader and more functional insight into the microbial community, particularly when reference genomes are lacking. Oral rinse sampling is a practical, non-invasive approach suitable for large-scale studies, enabling effective preservation and transport of DNA^22,25,33^.

The aim of the study was to analyze microbiome of healthy human oral cavity and to determine *in vitro* antimicrobial properties of A-PRF (platelet-rich fibrin in the form of a membrane obtained after blood centrifugation) and LP (the liquid fraction of plasma remaining after centrifugation) obtained from people with characterized during our study microbiome, against Gram-negative bacteria: *E. coli, P. gingivalis* and Gram-positive bacteria creating different shapes and forms: *Bacillus subtilis*, *Micrococcus luteus*, *Staphylococcus lentus*, *Enterococcus casseliflavus*, and *Streptococcus mutans*.

## 2. Results

### 2.1. Study participants

All study participants were in good health, were nonsmokers, had no symptoms of oral infection, possessed their natural dentition and had taken no antibiotics for at least 1 month prior to the experiments. Finally, 20 patients, including 9 men and 11 women were selected based on a conducted interview (Supplementary material 1: ‘Patient survey’ and ‘Patient examination form (API)’), blood test results (Supplementary material 2: Table S1.) and after an oral cavity examination.

### 2.2. Bacterial community composition

Bacterial community structure at the phylum is visualized in Fig. 1 and the core microbiome for the samples for phylum is shown in Fig. 2.

**Figure 1.**
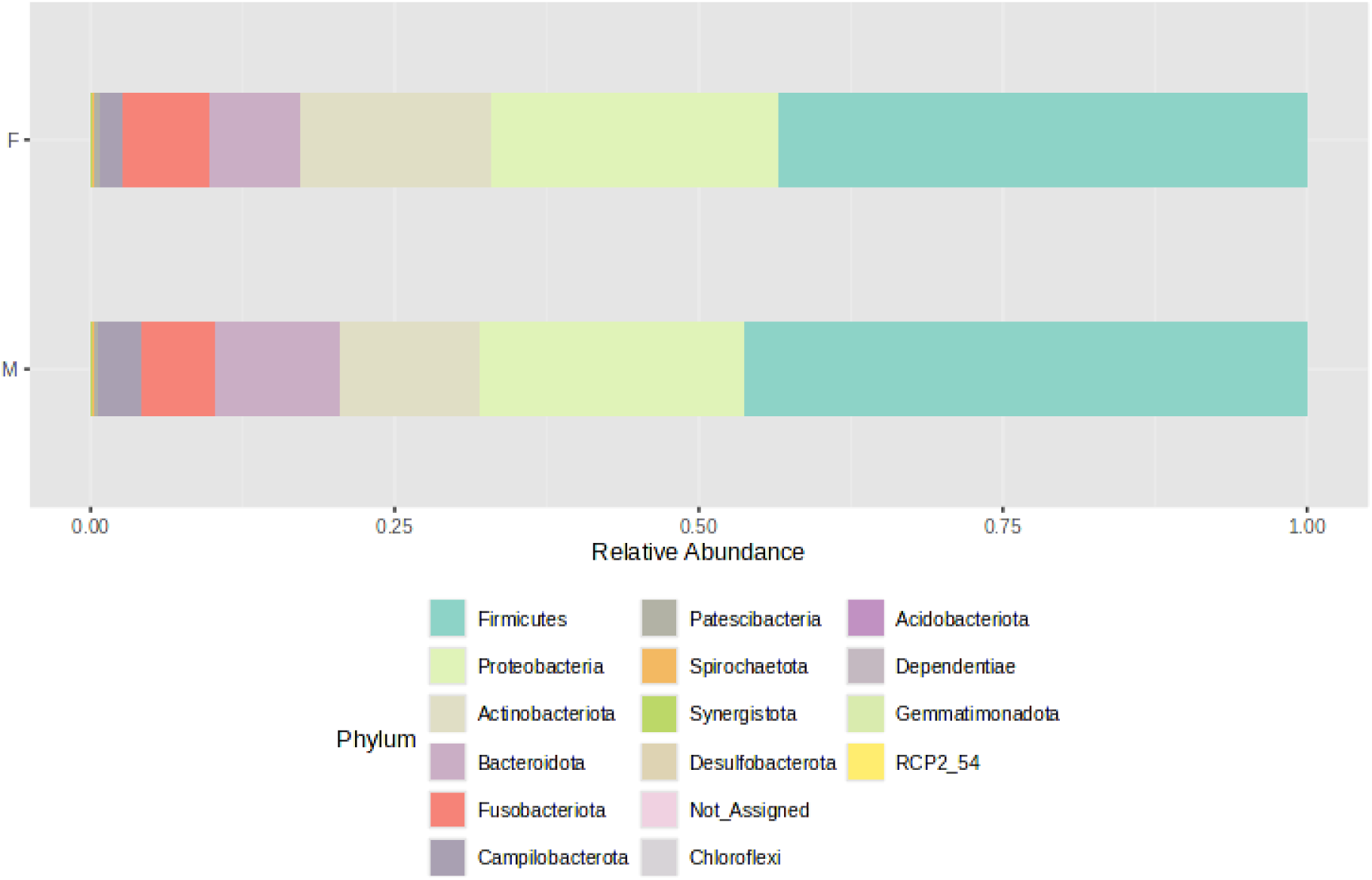
Bacterial community structure composition of samples at the phylum level. The samples were divided into two groups - F (female) and M (male); stacked barplots represents average value for all samples assigned to groups.

**Figure 2.**
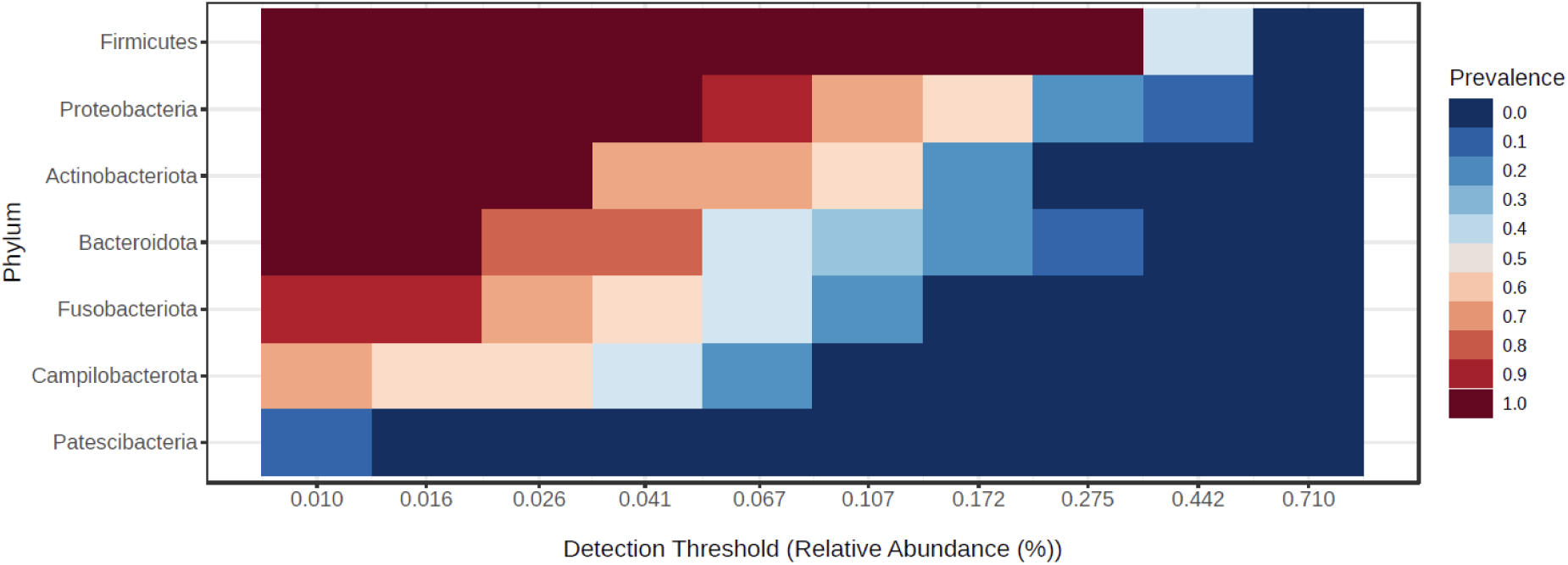
Core bacteria phyla across all samples.

For all samples originated from female and male, the alpha-biodiversity indexes have been calculated: Chao1 (estimates species richness, emphasizing rare species), Shannon (accounts for both richness and evenness), Simpson (measures dominance) and Pielou (quantifies the evenness of species distribution within a sample), and they do not differ between both analyzed groups, namely female and male (p>0.05). Moreover, beta-diversity has been analyzed and the Bray-Curtis index, which quantifies community dissimilarity based on species abundance (considering both shared and unique species between samples) and the Jaccard index, which measures compositional similarity, focusing solely on species presence or absence without accounting for abundance do not differ between female and male samples (p>0.05). Boxplots for diversity indexes data distribution are presented in Supplementary material 3: Microbiome biodiversity indexes comparison between female and male patients.

Microbiome analyses (Fig. 1) revealed that the majority of bacterial phyla belonged to the *Firmicutes*, followed by *Proteobacteria* and *Actinobacteria*, with lower relative abundances of *Bacteroidota*, *Fusobacteriota*, and *Campylobacteriota*. The dominant bacterial classes were *Bacilli*, *Gammaproteobacteria*, *Actinobacteria*, *Negativicutes*, and *Bacteroidia*, followed by *Fusobacteria*, *Campylobacteria*, and *Clostridia*, listed in order of relative abundance. This taxonomic distribution pattern was consistent across samples obtained from both male and female participants. These findings are in line with diversity index analyses, which indicated no significant differences in microbial diversity between the groups.

The core microbiome analysis (Fig. 2) demonstrated that *Firmicutes* represent the dominant and most consistently present phylum across samples, indicating their role as a key component of the core microbiome. *Proteobacteria* were also highly prevalent, though generally detected at lower relative abundances than *Firmicutes*. *Actinobacteriota* appeared to be a relatively stable, albeit less abundant, member of the microbiome. In contrast, *Bacteroidota* were commonly detected but rarely exceeded higher abundance thresholds. *Fusobacteriota*, *Campylobacterota*, and *Patescibacteria* were found infrequently and at low abundance levels, suggesting they may constitute part of the variable or transient microbiome. Notably, only *Firmicutes*, *Proteobacteria*, and *Actinobacteriota* maintained both high prevalence and moderate abundance, supporting their classification as core phyla.

The correlation network between bacterial classes identified in the oral cavity (Fig. 3) reveals patterns of microbial co-occurrence and potential ecological interactions. Each node represents a bacterial class, with color indicating the origin of the sample: orange for female-derived samples, green for male-derived samples, and bi-colored nodes representing taxa present in both types of samples. Edges indicate statistically significant correlations in relative abundance, suggesting co-occurrence. A substantial number of bacterial classes, including *Clostridia, Bacteroidia, Negativicutes, Gammaproteobacteria, Fusobacteriia*, and *Bacilli*, appear in both male and female samples and form a dense central cluster. This suggests the presence of a core microbial community that is shared across sexes and potentially functionally integrated through ecological interactions. The dense connectivity of these bacterial classes may indicate synergistic or co-dependent metabolic relationships within the oral microbiome. In contrast, certain bacterial classes such as *Desulfotobacteria, Babeliae*, and the unclassified RCP2_54 appear to be specific to female samples, while *Acidimicrobia, Longimicrobia*, and *Chloroflexia* are uniquely associated with male samples. These findings suggest the existence of sex-specific components of the oral microbiota; however, the analysis of biodiversity indexes does not confirm these findings. Oral microbiome may be additionally influenced by factors such as hormonal environment, behavioral habits, or differential exposure to environmental factors.

**Figure 3.**
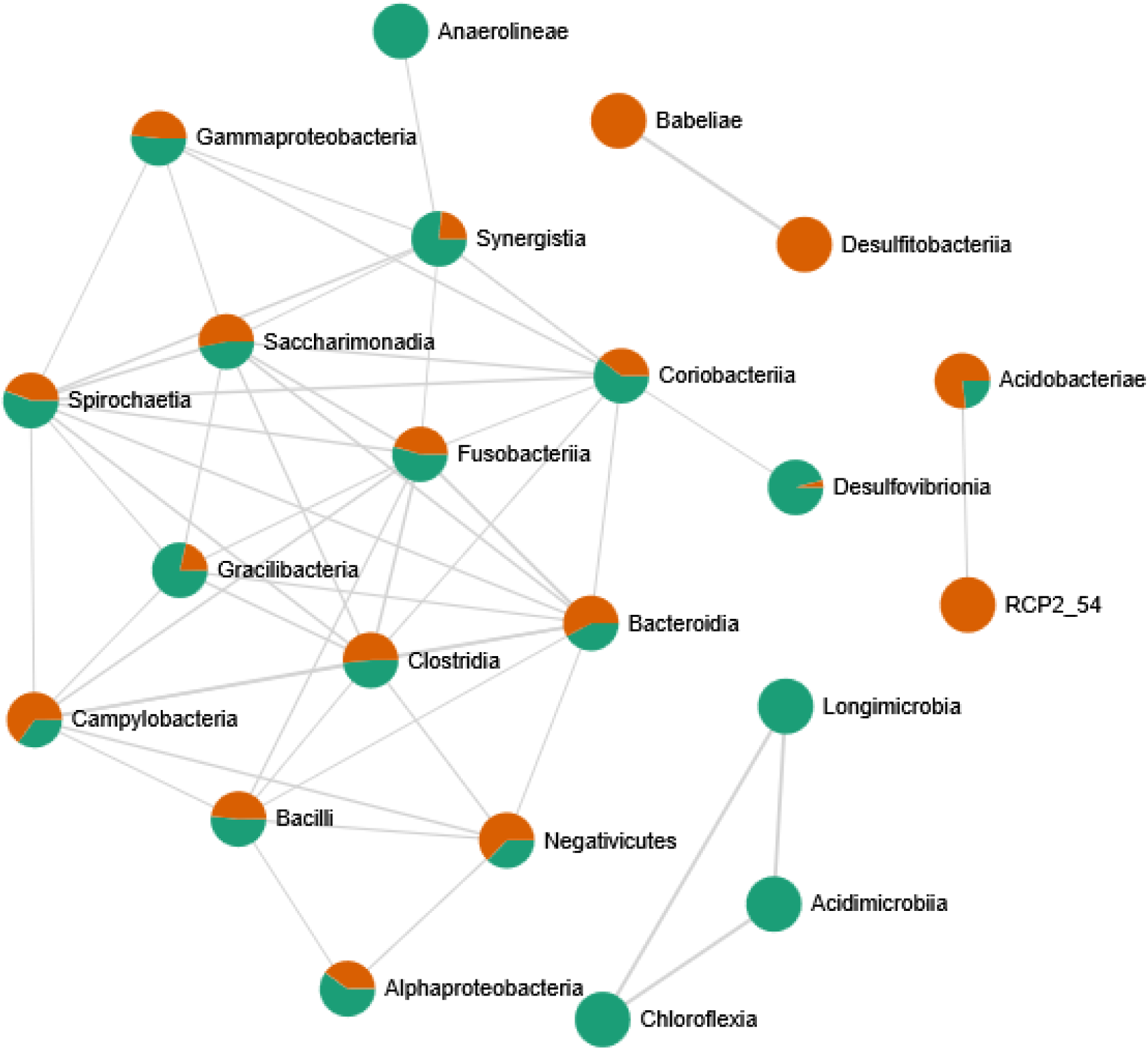
Correlation network between bacterial classes identified in the oral cavity. Network was created by the MicrobiomeAnalyst web tool. Only strong (|r|>0.5) and significant (p<0.05) correlations are presented. Nodes as a pie chart represents classes’ relative abundances. The colour of the nodes indicates: female samples (green) and male samples (orange)

### 2.3. Fungal community structure

Fungal community structure at the phylum level is visualized in Fig. 4 and the core mycobiome for the samples for phylum levels are shown in Fig. 5.

**Figure 4.**
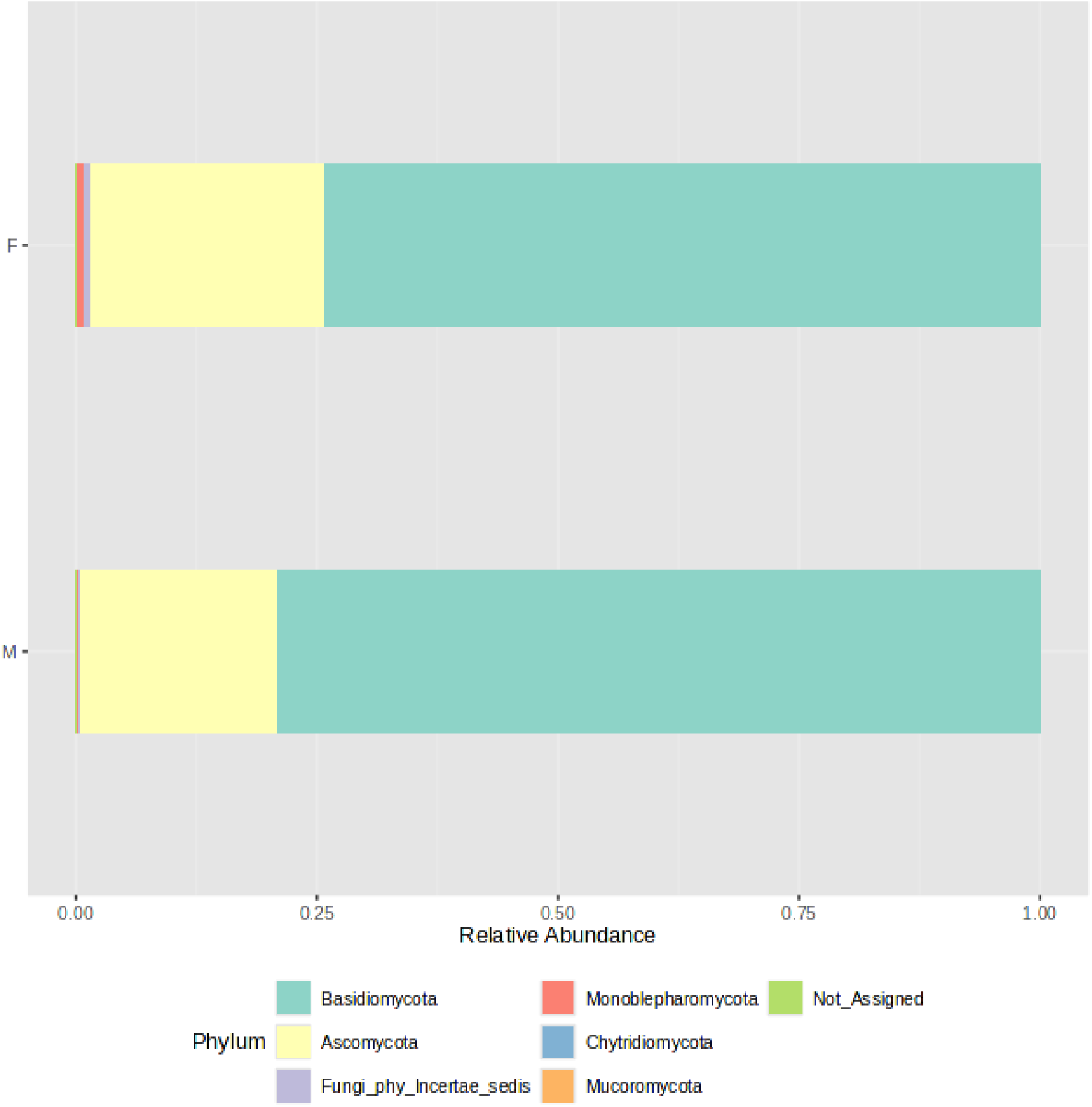
Fungal community structure composition of samples at the phylum level. The samples were divided into two groups - F (female) and M (male); stacked barplots represents average value for all samples assigned to groups.

**Figure 5.**
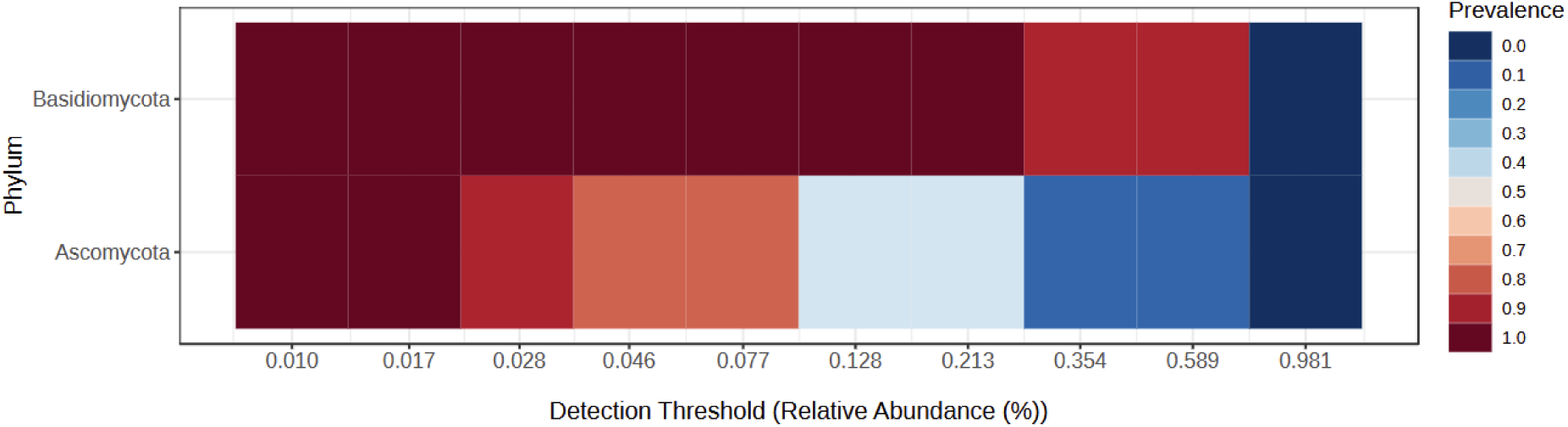
Core fungal phyla across all samples.

For all samples the alpha-biodiversity indexes have been calculated: Chao1 (estimates species richness, emphasizing rare species), Shannon (accounts for both richness and evenness), Simpson (measures dominance) and Pielou (quantifies the evenness of species distribution within a sample), and they do not differ between both analysed groups, namely Female and Male (p>0.05). Furthermore, beta-diversity has been analysed and the Bray-Curtis index and the Jaccard index do not differ between groups (p>0.05). Boxplots for diversity indexes data distribution are presented in Supplementary material 3: Microbiome biodiversity indexes comparison between female and male patients.

Fungal community composition (Fig. 4) in oral samples shows a dominance of two main phyla: *Basidiomycota* and *Ascomycota*. In both male and female samples, *Basidiomycota* represents the most abundant phylum, accounting for approximately three-quarters of the total fungal community. *Ascomycota* forms the second most prominent phylum, constituting about one-fourth of the total relative abundance. Other fungal phyla such as *Mucoromycota*, *Chytridiomycota*, and *Monoblepharomycota* are present in much lower proportions. Their detection suggests a minor and potentially transient role in the oral ecosystem. A small fraction of the sequences remains unassigned (Not_Assigned) at the phylum level, which may indicate the presence of novel or less-characterized fungal phyla. Importantly, the overall fungal composition is highly similar between sexes, with no major differences in the relative proportions of dominant phyla. This suggests a relatively conserved fungal core microbiome across male and female individuals in the studied population. This was also confirmed by analysis of different biodiversity indexes (Supplementary material 3: Microbiome biodiversity indexes comparison between female and male patients).

The core microbiome analysis (Fig. 5) demonstrates that *Basidiomycota* exhibits consistently high prevalence across nearly all detection thresholds of relative abundance. This indicates that members of this phylum are both widely distributed and consistently present in the majority of samples, making *Basidiomycota* a clear component of the core oral mycobiome. In contrast, *Ascomycota* shows a broader range of prevalence, with high detection at lower abundance thresholds, but decreasing prevalence at higher thresholds. This suggests that while *Ascomycota* is also a frequent constituent of the oral fungal community, it may exhibit greater variability in abundance among individuals, potentially reflecting individual differences or transient colonization. Overall, this heatmap supports the conclusion that *Basidiomycota* forms the most stable and dominant component of the core oral mycobiome, while *Ascomycota* may represent a more variable, yet common, element of the community.

The fungal co-occurrence network (Fig. 6) illustrates associations between different fungal classes identified in oral samples, with nodes colored according to their relative representation in male and female subjects. Mixed-colored nodes indicate classes present in both sexes, albeit at varying abundances. *Saccharomycetes* appears as a central and well-connected node, forming links with *Taphrinomycetes* and *Malasseziomycetes*, suggesting potential co-occurrence patterns or shared ecological niches in the oral environment. These classes are commonly found in both male and female samples and likely represent stable components of the core mycobiome. A separate cluster includes *Cystobasidiomycetes* and *Fungi_cls_Incertae_sedis*, both predominantly found in male samples. This may indicate sex-specific factors influencing the presence or abundance of these fungi, such as hormonal differences, lifestyle, or oral hygiene habits. Other classes, such as *Eurotiomycetes* and *Dothideomycetes*, show a slightly higher prevalence in female samples. Meanwhile, a distinct group composed of *Pucciniomycetes*, *Spizellomycetes*, and *Dacrymycetes* appears isolated and is found exclusively or primarily in male samples, possibly reflecting transient or niche-specific colonizers. The analysis of different biodiversity indexes does not confirm these assumptions (Supplementary material 3: Microbiome biodiversity indexes comparison between female and male patients). Overall, the network highlights the existence of a core group of fungal classes common across individuals, as well as variable or rare taxa that may reflect individual-specific or environmental factors. The patterns of co-occurrence provide insights into the complexity of the oral mycobiome.

**Figure 6.**
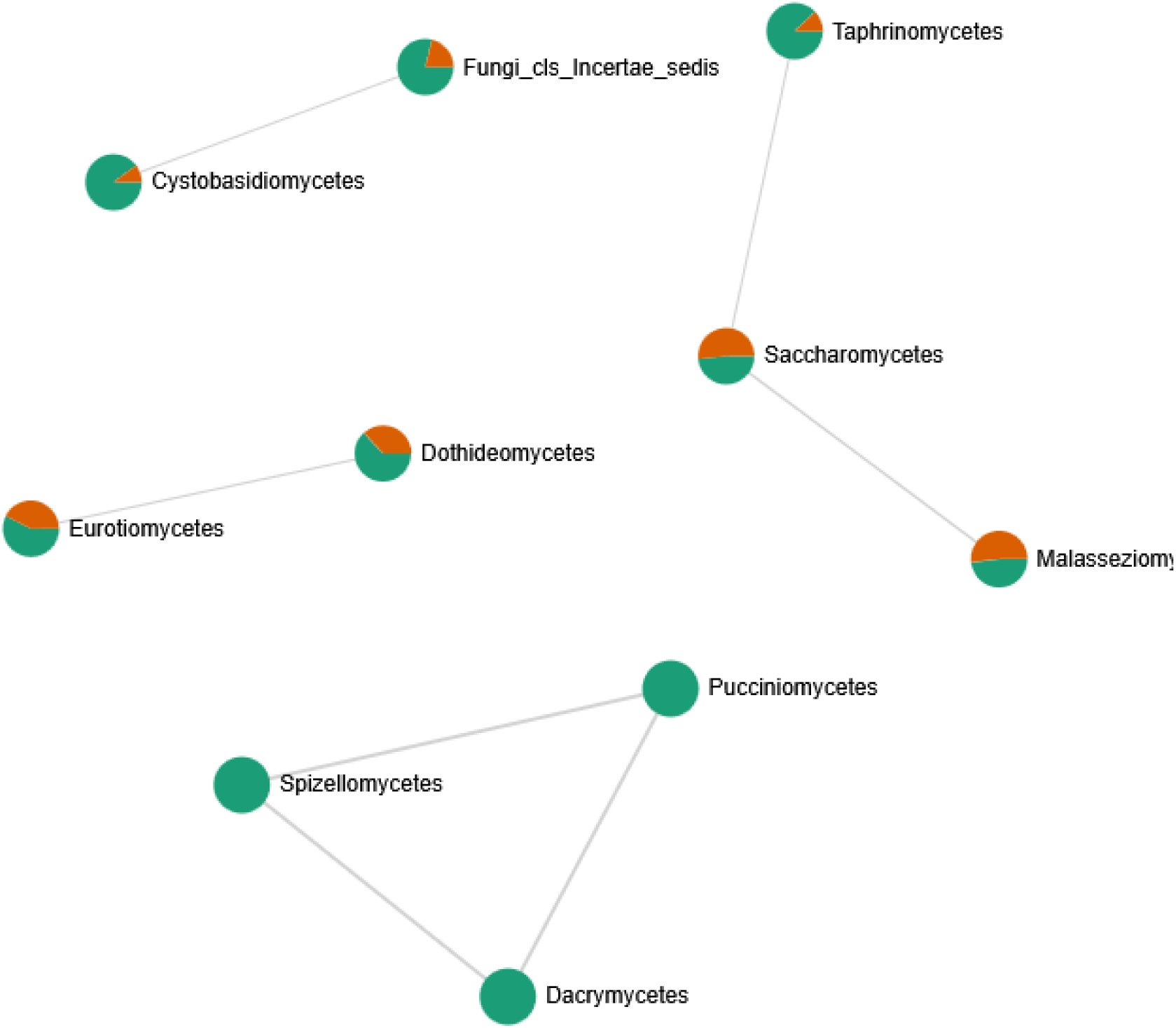
Correlation network between fungal classes identified in the oral cavity. Network was created by the MicrobiomeAnalyst web tool. Only strong (|r|>0.5) and significant (p<0.05) correlations are presented. Nodes as a pie chart represents classes’ relative abundances. The colour of the nodes indicates: female samples (green) and male samples (orange)

### 3.4. Antibacterial effect of the A-PRF membranes and liquid fraction LP

The antibacterial activities of the A-PRF membrane and LP fraction were analyzed by directly placing them on the surface of a medium covered in bacterial suspension. Activity against bacteria with different cell wall structures and also having different shapes and creating different structures was studied. In all tested systems, after the defined incubation period and conditions, antimicrobial activity was demonstrated as a clear zone around the A-PRF membrane, the cellulose discs soaked with the LP fraction and the LP fraction dropped on the surface of medium covered in bacterial suspension. In each case, a wider inhibition zone was observed for *E. coli* (average 4 mm) compared to the tested Gram-positive bacteria (0-3 mm) - for *E. casseliflavus* - 0, *M. luteus* and *B. subtilis* - 1 mm, *S. lentus* - 3 mm. Interestingly, the clear zones were better visible after 24 h of incubation than after 48 h, where overgrowing of this zone with bacteria was observed in each case. Anaerobic bacteria: *S. mutans* and *P. gingivalis* were found to have the lowest sensitivity, the growth inhibition zone was less than 1 mm after 24 h and 48 h, respectively, but after further incubation, this zone became completely overgrown.

For the analysis of bacterial growth in a liquid medium in the presence of A-PRF membrane or LP fraction, a growth inhibition effect was observed after 6 h for all bacteria compared to the control culture, but the strongest effect was noted for *E. coli* (Fig. 7). Interestingly, for Gram-positive bacteria, the least sensitive were cocci forming structures and capable of forming biofilms: *E. casseliflavus*, *M. luteus*, and *B. subtilis,* but *S. lentus*, although also capable of forming a biofilm, was definitely more sensitive than other Gram-positive bacteria tested (Fig. 7). The highest differences in OD_600_ readings compared to the control culture was observed after 2 h and 4 h of incubation. Antibacterial activity was also confirmed by a lower CFU/ml value at selected time points (Table 1, group A). We also evaluated the condition/size of fragmented A-PRF (titration plates) and a whole A-PRF membrane (liquid culture - 3 ml) after 24 h of incubation with aerobic bacterial cultures. All analyzed membranes remained unchanged.

**Figure 7.**
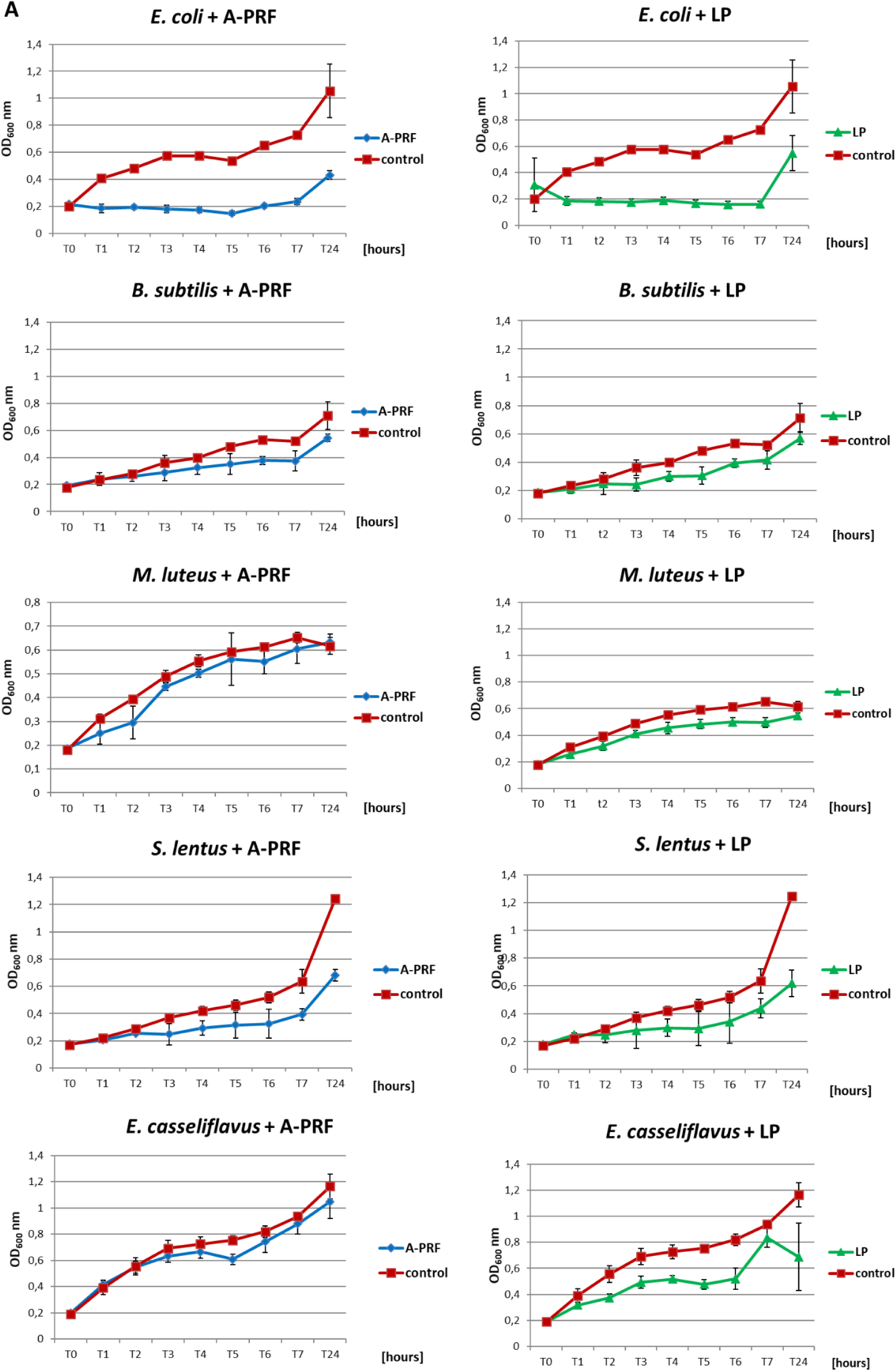
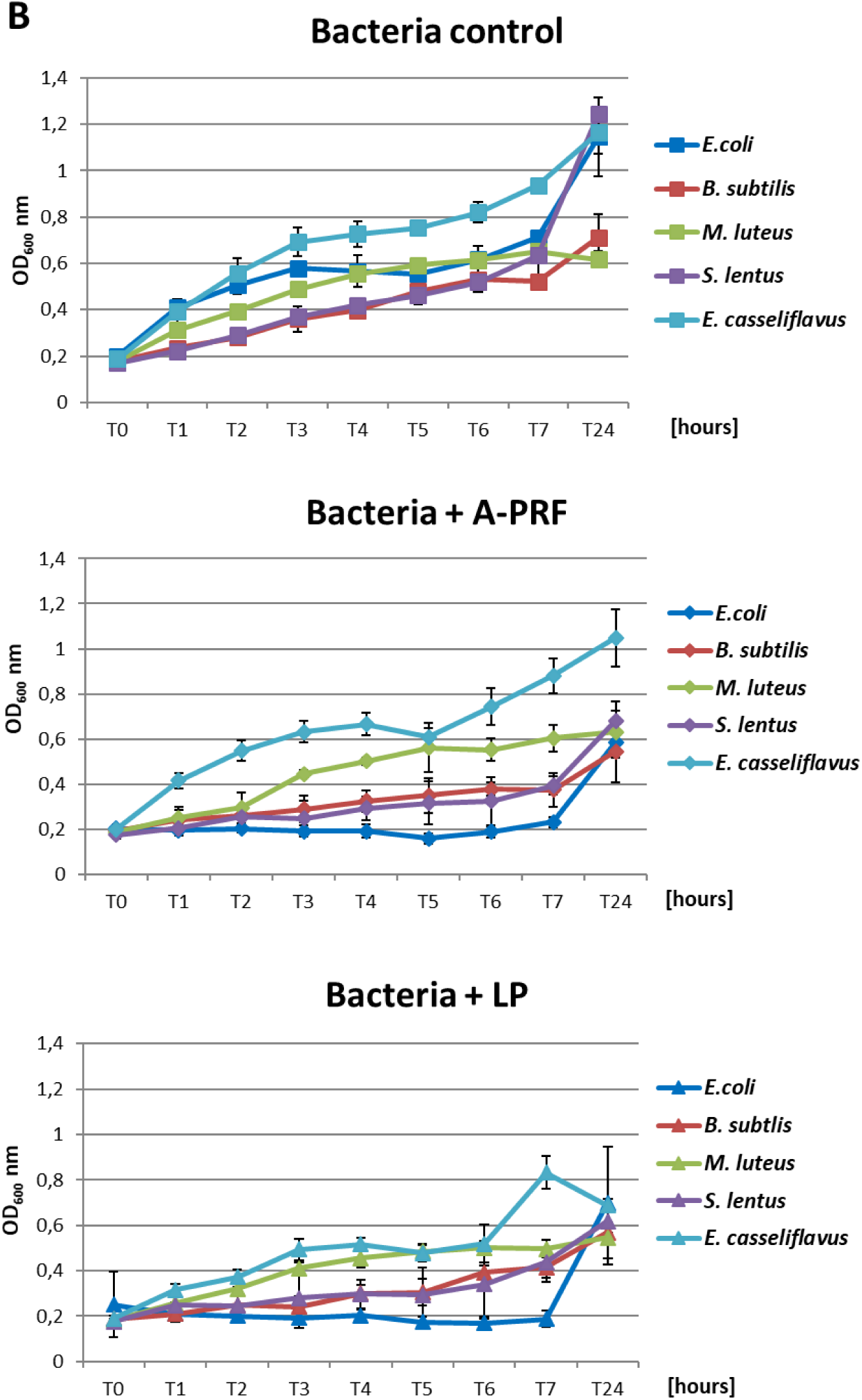
The growth curves of the tested bacteria from group A: *E. coli, B. subtilis, M. luteus, S. lentus, E. casseliflavus* in liquid medium (titration plates), in the presence of A-PRF or LP fraction, incubated at 37 ℃ in aerobic conditions. **A** - Individual charts; **B** - collective charts

**Table 1.**
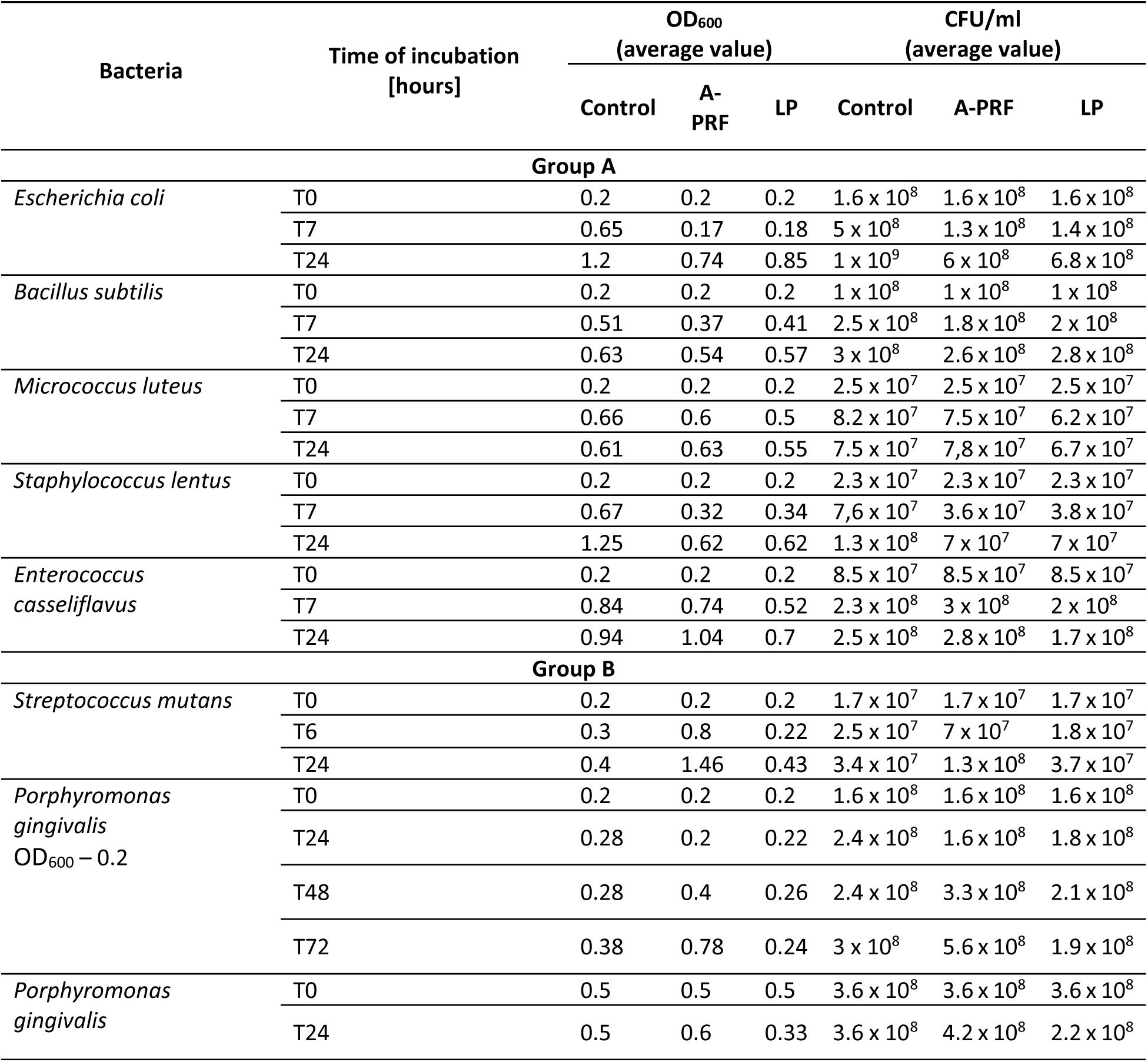
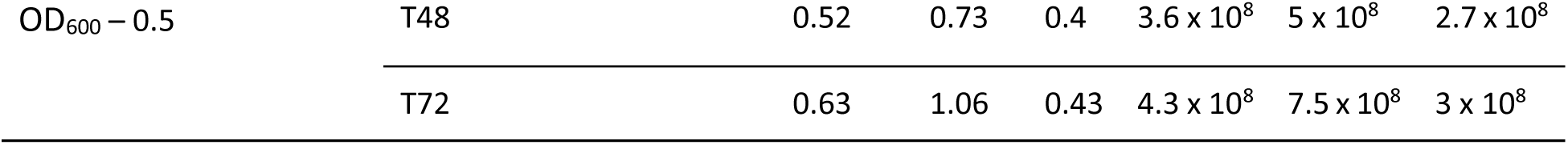
The values of OD_600_ and CFU/ml obtained in the tests for group A and B bacteria, taking into account the incubation time.

In the case of *P. gingivalis*, where the OD_600_ at time point T0 was 0.2, a slight inhibition of bacterial growth was observed after 6 h with the A-PRF membrane: the OD_600_ increased by an average of 1.5-fold, whereas in the control sample (without A-PRF), the OD_600_ increased nearly threefold. In a parallel experiment with a higher initial OD_600_ value of 0.5 at T0, a moderate increase in OD_600_ was recorded in the culture supplemented with the A-PRF membrane (2.5-fold), compared to the control, where OD_600_ increased by an average of 1.7-fold. After 24 hours, OD_600_ values of both culture variants converged. For *S. mutans*, no significant differences in OD_600_ values were observed between the experimental and control conditions after both 6 h and 24 h of incubation (Table 1, group B). Due to the generation time of *S. mutans* of 2 hours and *P. gingivalis* of an average of 7 hours, it was decided to perform an analogous study using 48-well titration plates and OD_600_ for *S. mutans* at T0, T2, T4, T6 and T24 and for *P. gingivalis* at T0, T24h, T48h and T72h. For *P. gingivalis*, with an initial OD_600_ around 0.2, an antibacterial effect of the A-PRF membrane was observed after 24 h of incubation. However, this effect was no longer evident after 48 h, as an increase in OD_600_ compared to the control culture was observed. After 72 h of incubation, the optical density had nearly doubled relative to the control. The application of the LP fraction had minimal impact on the bacterial growth level (Fig. 8A). When the initial OD_600_ was approximately 0.5, no antibacterial effect of the A-PRF membrane was detected, and after 72 h, a more than 1.5-fold increase in OD_600_ compared to the control was observed. In the case of the LP fraction, a slight decrease in OD_600_ was noted only after 24 h of incubation (Fig. 8B). The results obtained in the study using *S. mutans*, with an initial optical density of about 0.2, revealed a lack of sensitivity to the antibacterial activity of the A-PRF membrane and even observed growth stimulation. After 6 h of incubation, there was a more than 3-fold increase in OD_600_ and after 24 h as much as 4.5-fold in comparison with the control culture (Fig. 8C). The use of the LP fraction had no effect on the level of growth of this bacteria (Fig. 8). Antibacterial activities or their lack was also confirmed by a CFU/ml value (Table 1, group B). Interestingly, after 24 h of incubation, the A-PRF membrane was completely dissolved in cultures of both bacterial species, both in the case of whole membranes (liquid culture - 3 ml) and their fragments (titration plates).

**Figure 8.**
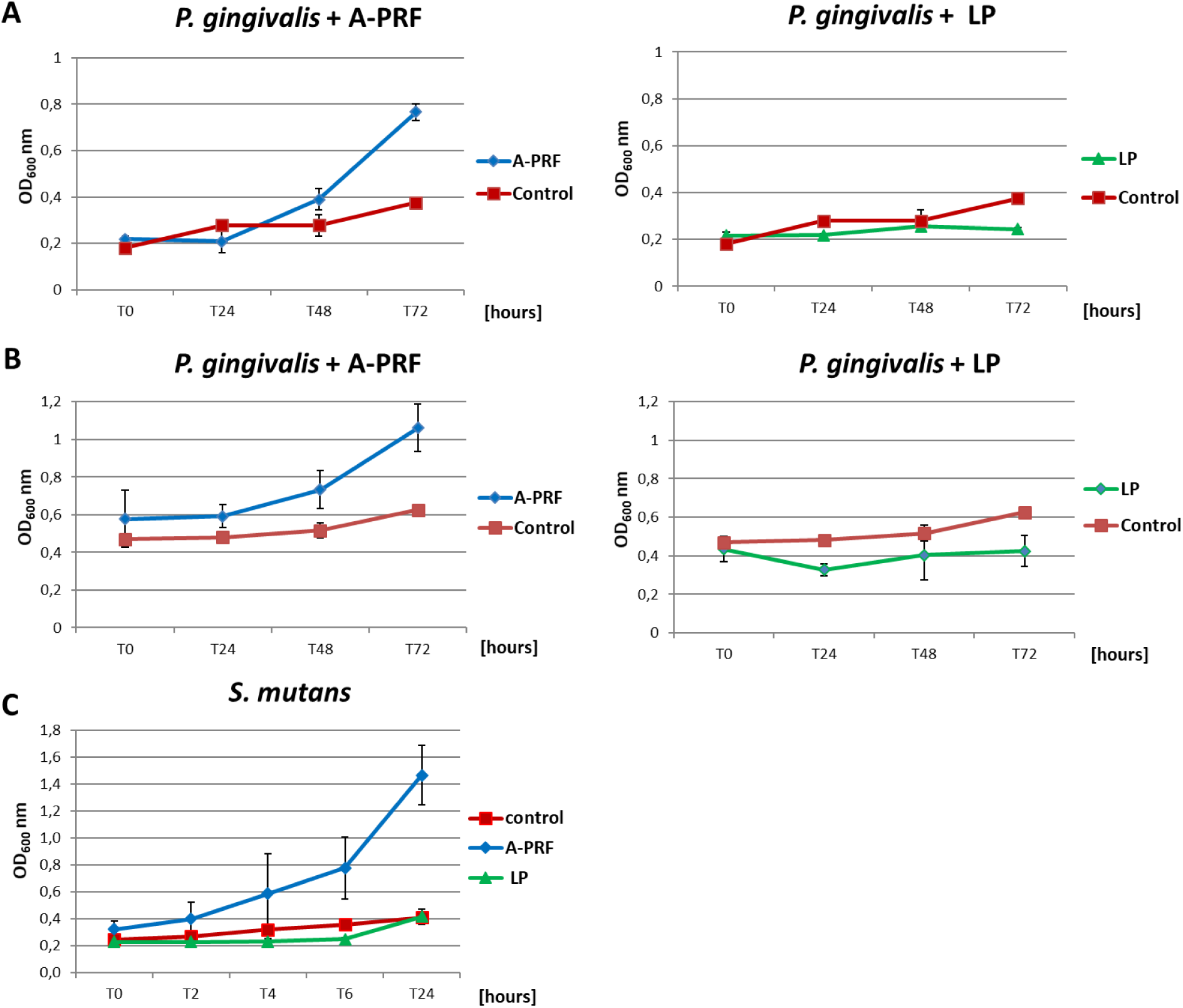
The growth curves of the tested bacteria from group B: *P. gingivalis* and *S. mutans* in liquid medium, in the presence of A-PRF or LP fraction, incubated at 37 ℃ in anaerobic conditions. **A** - *P. gingivalis* with an initial OD_600_ approx. 0.2; **B** - with an initial OD_600_ approx. 0.5; **C** - *S. mutans*

## 3. Discussion

Antibacterial resistance is currently recognized as a serious global health threat. Due to misuse or overuse of antibiotics and other medications, pathogens are no longer susceptible to available treatment that leads to disease spread and death of patients. WHO estimates that AMR contributed to 4.95 million deaths in 2019^34^. The primary action to address this problem is the search for new antimicrobial substances or the modification of previously described concepts/resolutions. An example of such an approach is the use of advanced platelet-rich fibrin, described in this work. Our study design was comprehensive, consisting of clearly defined and logically sequenced stages. Patient qualification involved an evaluation of survey responses, blood test results, and clinical examination of the oral cavity. The study enrolled systemically healthy, non-smoking individuals with no signs of oral infection, a complete natural dentition, and no antibiotic use for at least one month prior to sampling. Subsequently, oral microbiome analysis was performed on the selected individuals. After confirming that biodiversity indices did not differ significantly between males and females, the entire cohort was considered a homogeneous study group.

Zaura et al.^35^ identified *Firmicutes* (including *Streptococcus* spp., *Granulicatella* spp., and members of the *Veillonellaceae* family), *Proteobacteria* (*Neisseria* spp., *Haemophilus* spp.), *Actinobacteria* (*Corynebacterium* spp., *Rothia* spp., *Actinomyces* spp.), *Bacteroidetes* (*Prevotella* spp., *Capnocytophaga* spp., *Porphyromonas* spp.), and *Fusobacteria* (*Fusobacterium* spp.) as the predominant taxa in the oral microbiome. Despite analyzing only three healthy individuals with differing geographic origins and ages, their study revealed significant inter-individual variability. Notably, their assessment of oral health was based solely on clinical periodontal status without consideration of biochemical or immunological markers. Despite methodological and demographic differences, our findings were largely consistent with theirs at the phylum level.

Our results partially aligned with those reported by Verma et al.^24^ and Nearing et al.^25^, who identified *Actinobacteria*, *Fusobacteria*, *Proteobacteria*, *Firmicutes*, and *Bacteroidetes*, respectively, as the most prevalent oral phyla. In contrast, we observed *Firmicutes*, *Proteobacteria*, *Actinobacteria*, *Bacteroidota*, and *Fusobacteria* as the dominant phyla. Nevertheless, mentioned studies consistently reported that these five phyla constitute approximately 90% of the oral microbial community.

In the present study, we analyzed 20 samples obtained from healthy male and female participants. Although the data were initially treated as a single cohort, we also investigated potential sex-based differences in microbial composition. No differences were observed in alpha- or beta-diversity indexes, likely due to high intra-sample variability or the influence of uncontrolled variables (not included in the participant survey). However, co-occurrence network analysis revealed microbial associations suggestive of sex-specific structuring. These findings imply that additional factors—such as hormonal status, behavioral differences, or environmental exposures—may contribute to shaping the oral microbiota in a sex-dependent manner. For example, hormonal fluctuations during the menstrual cycle are associated with changes in the relative abundances of *Campylobacter* spp., *Haemophilus* spp., *Oribacterium* spp., and *Prevotella* spp.^36^, and pregnancy has been linked to increased levels of *Neisseria* spp., *Porphyromonas* spp., and *Treponema* spp., while *Streptococcus* spp. and *Veillonella* spp. are more abundant in non-pregnant women^37^.

Furthermore, the oral microbiome has been implicated in the pathogenesis of systemic diseases, including cancers (e.g., head and neck, pancreatic, colorectal), as well as autoimmune and neurodegenerative disorders such as rheumatoid arthritis, hypertension, Alzheimer’s disease, and systemic lupus erythematosus^31,38^. Dietary patterns also modulate the oral microbiota: intake of fiber, medium-chain fatty acids, piscine monounsaturated fatty acids, and polyunsaturated fatty acids is associated with increased microbial diversity, while consumption of sugar and refined carbohydrates correlates with higher abundances of specific bacterial taxa. Notably, carbonated beverage consumption is positively associated with the presence of *Bacteroidetes*, *Gammaproteobacteria*, *Fusobacterium*, and *Veillonella*^39,40^.

Environmental factors such as smoking are known to significantly alter oral microbial composition, particularly by favoring anaerobic taxa^41^. Oral hygiene practices also play a crucial role in shaping microbial diversity^42^. Geographic and climatic differences can further modulate the oral microbiome, potentially contributing to inter-individual variation at the species and strain levels^38,41^. However, in the present study, we controlled for these two specific factors by recruiting only non-smokers residing in the same geographic area.

Despite numerous individual-specific influences on the oral microbiota, core microbiome analysis can serve as a reference point for microbial eubiosis. The oral core microbiome, defined as the set of consistently shared taxa and their genetic content across individuals, reflects a functionally important microbial backbone. Caselli et al.^43^ demonstrated a lack of statistically significant differences in core microbiome composition among study participants, supporting its potential utility as a benchmark for oral health.

In our dataset, core microbiome analysis identified *Firmicutes*, *Proteobacteria*, and *Actinobacteriota* as the most consistently prevalent and moderately abundant phyla, reinforcing their designation as key components of the oral core microbiota. In contrast, *Bacteroidota*, *Fusobacteriota*, *Campylobacterota*, and *Patescibacteria* were detected less consistently or at lower relative abundances, suggesting a more variable or transient role. Taxa found predominantly at low abundance thresholds may represent niche-specific or functionally specialized community members. This analytical approach thus enables the differentiation between stable, shared microbial constituents and more labile taxa influenced by host-specific or environmental variables. Our findings were partially consistent with previous reports^2,24^, identifying similar core phyla but differing in relative abundance and detection thresholds.

It has been found that fungi account for 0.004% (approximately 100 species)^28,44^ of the overall oral microorganisms and have been detected in specimens from the hard palate, supragingival plaque, and oral rinses^43^, but also in saliva samples (*Malassezia* spp. and *Candida* spp.)^38^. Due to the lack of data aiming at a healthy human oral mycobiome, it is hard to conduct comprehensive analysis, but we found *Basidiomycota* and *Ascomycota* representing three-quarters of the total fungal community. *Basidiomycota* is commonly associated with environmental fungi, but some members may also be part of the normal oral microbiota^45^, and *Ascomycota* include various yeasts and filamentous fungi, some of which are frequently found in the human oral cavity e.g., *Candida* spp. The common fungi genera found in the oral cavity together with *Candida* spp. are *Aspergillus* spp.*, Aureobasidium*s spp.*, Cladosporium* spp.*, Cryptococcus* spp.*, Fusarium* spp.*, Gibberella* spp.*, Penicillium* spp.*, Rhodotorula* spp.*, Saccharomycetales* spp.*, and Schizophyllum* spp.^28,44^. Moreover, *Candida* spp. is widely reported as a part of a healthy human oral microbiome, but also during disease^38^.

Given the absence of differences in microbiome diversity indices between sexes, all samples were pooled into one dataset, and antimicrobial activity of obtained PRF were assessed. To date, antibacterial activity of PRF has been demonstrated against Gram-negative bacteria: *E. coli*, *Proteus mirabilis* and *Pseudomonas aeruginosa* and Gram-positive bacteria: *Bacillus megaterium*, *Enterococcus faecalis* or *S. aureus* and *S. epidermidis*^9,11,46^. It is worth emphasizing that the few *in vitro* studies on the antibacterial properties of PRF were conducted on a small scale and usually using one method: analysis of growth inhibition zones around PRF using the agar diffusion technique or counting CFU/ml after the bacterial cells incubation with PRF and only after one specific incubation time. It is therefore difficult to refer to these findings in the context of the results presented in this publication. Feng et al.^11^ demonstrated antimicrobial activity of PRF against *S. aureus* and *E. coli*, using the agar-based diffusion assay (incubation time with PRF - 24 h) and plate-counting test methods (spread on plates after 4 hours of incubation of bacteria with PRF). Research conducted by Jasmine et al.^9^ allowed the determination of minimal inhibitory concentration (MIC) and minimal bactericidal concentration (MBC) for the i-PRF fraction for *S. aureus* and *S. epidermidis* isolated from patients with oral and dental abscess after incubation at 37 ℃ for 24 h, using 96-well polystyrene microtiter plates and tryptone soya broth (TSB). It was also shown that i-PRF actively inhibited the biofilm formations of tested bacterial strains at 24 h^9^. In previous work, antibacterial activity was proven against: *E. coli, B. megaterium, P. aeruginosa, E. faecalis and P. mirabilis* by measuring inhibition zones in agar diffusion tests. Further studies were directed at characterizing the molecular mechanism responsible for the antibacterial activity of PRF against Gram-negative pathogens. These analyses confirmed the presence of hBD-2 (human beta-defensin 2), a known antimicrobial peptide^46^. A recently published study demonstrated antimicrobial effects of clindamycin-loaded PRF against *S. aureus, Streptococcus pneumoniae, Streptococcus mitis, P. gingivalis*, and *Fusobacterium nucleatum* using the agar-based diffusion assay^47^.

To date, no attempts have been described elucidating the reasons for higher antibacterial activity of PRF against Gram-negative bacteria compared to Gram-positive ones. Moreover, we did not find previous studies aiming at tracking the dynamics of changes over time of this activity, nor assessing the effects at varying initial bacterial concentrations. In the case of Gram-positive bacteria, the potential effect of PRF on different structural or morphological bacterial forms has also not been assessed. In our study, we addressed these gaps by designing experiments that account for these variables, thus providing a more comprehensive understanding of PRF’s antimicrobial potential. In our research we used: Gram-negative bacteria: *E. coli* (rod-shaped bacteria commonly found in the gut of humans and warm-blooded animals, but some *E. coli* strains do cause different illnesses e.g.: diarrhea, urinary tract infections, pneumonia, and even sepsis)^48^ and *P. gingivalis* (Gram-negative, rod-shaped, strictly anaerobic bacteria, a keystone pathogen in chronic periodontitis)^49^, and Gram-positive bacteria creating different shapes: *B. subtilis* (rod-shaped can forming chains, spore-forming bacteria, used as a probiotic, model organism in biotechnology and medicine research, can cause various diseases in humans, including food poisoning, bacteremia, endocarditis, sepsis and pneumonia)^50^; *M. luteus* (spherical bacterium, cocci occurs in tetrads, spherical bacteria, cocci occurs in tetrads, occurs all over the skin, rarely causes disease, although in people with weakened immune system can cause serious infections)^51^; *S. lentus* (cocci, occurring singly, in pairs or tetrads, mainly an animal pathogen, however, it can colonize humans and cause a number of clinical symptoms)^52^; *E. casseliflavus* (spherical, ovoid-shaped bacteria, forming pairs or short chains, occasionally cause opportunistic infections)^53^; *S. mutans* (cocci, typically form pairs or chains, facultatively anaerobic bacteria related to the etiology and pathogenesis of dental caries)^54^. Gram-positive and gram-negative bacteria differ primarily in their cell wall structure. Our study demonstrated that PRF, regardless of its form, exhibits antibacterial activity, with the strongest effect observed against the Gram-negative bacteria *E. coli*. Among the tested Gram-positive bacteria, highest activity was noted against *S. lentus*.

Generally, Gram-positive bacteria possess a thick peptidoglycan layer, with many different proteins and polymers (e.g., teichoic acids), as their primary cell wall component^55^, while gram-negative bacteria have a thin peptidoglycan layer surrounded by an outer membrane containing lipopolysaccharide^56^. For small molecules, the cell wall of Gram-negative bacteria does not pose a barrier; it is also thinner, which results in a possibly faster antibacterial effect. In the case of Gram-positive bacteria—especially those forming various structures such as clusters, chains, or packets—the presence of multilayered peptidoglycan and additional polymers reduces the surface area available for interaction between the antibacterial molecule and the bacterial cell envelope. Additionally, bacteria from this group can produce capsules and modify the charge of their cell wall, making them less susceptible to antibacterial compounds^57–59^.

A particularly important finding of this study is the demonstrated lack of antibacterial activity of the A-PRF membrane against *S. mutans*, as well as the stimulation of growth observed for *P. gingivalis*. Notably, for both species, complete degradation of the A-PRF membrane was observed after 24 hours of incubation, in contrast to the other tested bacterial strains. Both *S. mutans* and *P. gingivalis* are known biofilm-forming species and are associated with oral health disorders. We hypothesize that bacterial proliferation was boosted by degraded the A-PRF, becoming additional source of food (cleavage by gingipains – peptides as a source of C, N …etc.), as both species are linked to periodontitis—a condition characterized by progressive destruction of the tissues supporting the teeth. *P. gingivalis* is an obligate anaerobic bacteria that inhabits the gingival pocket and belongs to the red complex^60^. Its presence in periodontal tissues has been identified in 78% of patients with periodontitis and only in 34% of healthy people^61^. The main source of carbon and energy for this bacteria are oligopeptides^62^. The most important virulence factors of *P. gingivalis* include fimbriae, hemolysin, hemagglutinin, capsules, outer membrane vesicles, lipopolysaccharides, and gingipains^63,64^. Gingipains are cysteine endopeptidases responsible for 85% of the proteolytic activity of this bacteria^65^. There are three types of gingipains distinguished by their specificity – lysine-specific gingipain Kgp and arginine-specific gingipain A (RgpA) and gingipain B (RgpB)^66^. These enzymes are present in all strains of *P. gingivalis* and play an important role in the physiology and virulence of this bacteria. *P. gingivalis* requires an exogenous source of heme for growth and in the subgingival pocket environment, red blood cells are a rich source of this substance. RgpA and Kgp have a hemagglutination-adhesion domain responsible for adhesion and penetration into erythrocytes, which consequently leads to cell disintegration, hemoglobin degradation and iron acquisition^67^. In addition, gingipains have the ability to bind to ECM (extra cellular matrix) components such as fibronectin, to degrade antibacterial peptides and to activate and degrade various elements of the immune system. These and many more gingipain activities result in the destruction of gingival tissue, alveolar bone loss as well as weakening of blood vessels^68^. The significant problem in the treatment of infections caused by this pathogen is the ability to form a biofilm. *P. gingivalis* in a biofilm is 500 to 1000 times less sensitive to antimicrobial drugs than planktonic cells^69^.

*S. mutans* possess several key virulence factors (e.g.: many different surface biological structures, surface proteins and adhesins) critically involved in the etiology and pathogenesis of dental caries. Through its ability to adhere to solid surfaces, *S. mutans* efficiently colonizes the oral cavity and initiates the formation of dental biofilm also known as dental plaque. This biofilm is characterized by a matrix of exopolysaccharides, which significantly influence its physical architecture and biochemical properties, thereby facilitating bacterial persistence^54^. Additional traits that enhance the colonization potential of *S. mutans* include its pronounced acidogenic capacity (acid production) and its ability to interact synergistically or antagonistically with other microbial species within the oral ecosystem^70^. Unlike the previously mentioned *P. gingivalis*, the main source of carbon and energy for this bacteria are carbohydrates. *S. mutans* can metabolize carbohydrates to produce acids that act on dental tissues to demineralize and form cavities^70^.

Our findings address existing knowledge gaps regarding the antibacterial activity of PRF against both pathogenic and opportunistic bacteria representing diverse taxonomic groups and morphological forms. These microorganisms are involved in a wide range of infections affecting the oral cavity, sinuses, and skin. By systematically analyzing the effects of PRF on bacteria with different cell wall structures and creating different forms, our study provides new insights into its spectrum of activity and highlights its potential clinical relevance.

## 3. Materials and methods

### 3.1. Participants and oral microbiome sample collection

The number of participants in the study was 30, including 15 men and 15 women, between 22 and 38 years old. Patients were selected based on a conducted interview (Supplementary material 1: ‘Patient survey’ and ‘Patient examination form (API)’), blood test results and after an oral cavity examination, which involved the assessment of oral hygiene using the Approximal Plaque Index (API). This index represents the percentage ratio of the number of interproximal tooth surfaces displaying biofilm to the total number of examined interproximal surfaces. Additionally, the presence of dental plaque was evaluated using a periodontal probe for interproximal surfaces from the buccal aspect in the first quarter of the dentition and from the palatal aspect in the second quarter. All participants were in good health, were nonsmokers, had no symptoms of oral infection, possessed their natural dentition and had taken no antibiotics for at least 1 month prior to the experiments.

The samples for DNA isolation were collected by swabbing for approximately 30 seconds the oral cavity, namely teeth, insides of surface of cheeks and gums, excluding tongue, using Epicentre Buccal Swab (2 swabs in each case). After sample collection, the swabs were placed in a transport system and moved in cooling conditions (4 ℃) to the laboratory. The isolation was done in sterile conditions at max. 30 minutes after samples were collected.

All the protocols used in this study were approved by the Ethics Committee of the Medical University of Warsaw (KB/94/2023).

### 3.2. DNA extraction

The total DNA was isolated from swabs using the commercially available kit for DNA purification from buccal swab samples (Swab; A&A Biotechnology, Gdynia, Poland) according to the manufacturer’s recommendation. The quantity and quality of extracted DNA were determined with a Qubit 4.0 Fluorometer (dsDNA high-sensitivity assay kit; Invitrogen, Thermo Fisher Scientific, Waltham, MA, USA) and a Colibri spectrophotometer (Titertek Berthold, Pforzheim, Germany), respectively. The DNA samples were isolated in triplicate and then pooled to obtain a single DNA sample for each sample. Only samples with concentrations higher than 10 ng/µL and an A260/A280 ratio ranging from 1.8 to 2.0 were analyzed. DNA samples were stored at −20 °C for further use.

### 3.3. Sequencing the variable V3–V4 regions of bacterial 16S rRNA

The structure of the bacterial community was determined by sequencing the variable V3–V4 regions of bacterial 16S rRNA (according to Illumina, Part # 15044223 Rev. B) and the fungal community structure was determined using the internal transcribed spacer ITS (ITS3/ITS4^71^) regions of fungal ribosomal DNA, both using the Illumina platform. Sequencing was performed on the Illumina NovaSeq 6000 instrument using the NovaSeq 6000 SP Reagent Kit v1.5 (500 cycles) in paired-end read mode (2×250 cycles), following the standard procedure recommended by the manufacturer with the addition of 1% PhiX control library. PCR and sequencing were done by the The CeNT’s Genomics Core Facility and the analysis were performed by the DNA Sequencing and Oligonucleotide Synthesis Facility (Institute of Biochemistry and Biophysics Polish Academy of Sciences). Raw sequences were processed and analyzed by QIIME2 software^72^ with the DADA2 option for sequence quality control and the newest release of the SILVA ribosomal RNA sequence database (SILVA SSU database 138.2) for taxonomy assignment^73,74^.

The amplicon data was visualized using the MicrobiomeAnalyst web server^75,76^. Dataset was not normalized, not scaled and not rarefied. The differences in the alpha-diversity in bacterial and fungal community structure were analysed using the QIIME2 pipeline, based on the Kruskal–Wallis H-test (Shannon, Chao1, Pielou and Simpson biodiversity indexes). The differences in beta-diversity were evaluated using the QIIME2 pipeline based on Bray-Curtis index. The correlations between different fungal phyla and different bacterial phyla were calculated and visualized using Spearman’s correlation coefficient with MicrobiomeAnalyst. The correlations between phyla were considered strong and significant when the absolute value of Spearman’s rank |r|> 0.9 and p < 0.05^77^.

### 3.4. Preparation of PRF fraction

From each participant, four venous blood tubes (A-PRF matrix sterile tubes, 10 ml, Dermoaroma Italy Srl 00144), each containing 8 ml, were collected and subsequently processed and centrifuged using a centrifuge (TD4C, Yingtai Instrument) for 14 minutes at 18.000 rpm at room temperature according to previous reports^19,78^. After centrifugation, the LP above the precipitated fibrin fraction was pipetted off. The collected LP was transferred to sterile 1.5 ml Eppendorf tubes. The A-PRF were compressed and converted into a standardized membrane with a thickness of 1 mm to determine their antibacterial abilities^79^. The clot containing the blood’s cellular components, located below, was removed. Two A-PRF membranes from each patient were used for further studies. To assess the participants’ hematopoietic system, an additional blood tube was collected for further peripheral blood morphology parameters analyzed in an external certified laboratory (Supplementary material 2: Table S1.).

### 3.5. Bacterial strains and general growth conditions

*P. gingivalis* ATCC33277 strain was grown in enriched tryptic soy broth (eTSB; composition per liter: 30 g trypticase soy broth, 5 g yeast extract, at pH 7.5; further supplemented with 5 mg hemin, 0.5 g L-cysteine, and 2 mg menadione) or on eTSB blood agar (eTSB medium containing 1.5% [wt/vol] of agar; further supplemented with 5% defibrinated sheep blood). *S. mutans* ATCC25175 strain was grown in eTSB. Both strains were incubated at 37 °C in anaerobic conditions with an atmosphere of 90% nitrogen, 5% carbon dioxide, and 5% hydrogen provided by Anoxomat® II Anaerobic Culture System^80^.

*E. coli, B. subtilis, M. luteus, S. lentus, E. casseliflavus* (collection of Institute of Microbiology, University of Warsaw) were grown in Luria-Bertani broth (LB) or on 1.5% LB agar plates at 37 °C in aerobic conditions, depending on experiment.

### 3.6. Antibacterial test of the A-PRF membranes and liquid fraction LP

To asses antimicrobial properties of A-PRF clot and LP using agar plates, cell suspensions (0.5 McFarland scale) of all strains: *E. coli*, *B. subtilis*, *M. luteus*, *S. lentus*, *E. casseliflavus, S. mutans* were spread evenly on Mueller Hinton II agar and *P. gingivalis* on eTSB blood agar with cotton swab (according to the Kirby-Bauer method^81^) and left aside for 15 minutes. Next, A-PRF clots were placed on a medium surface as a whole fragment or cut into smaller pieces (one whole A-PRF clot was cut on a maximum 5 smaller pieces of equal size)^11^. For LP analysis, sterile cellulose discs (Oxoid, Thermo Scientific) soaked with 20 µl of the LP fraction were placed on the surface of medium covered in bacterial suspensions. Additionally, LP liquid was also directly applied (20 µl) onto plates. Disks soaked with 20 µl of sterile phosphate buffered saline (PBS, ph 7.4; Merck, Germany) served as a negative control. Plates were incubated for 24h at 37 ℃ in aerobic conditions for *E. coli*, *B. subtilis*, *M. luteus*, *S. lentus*, *E. casseliflavus* (group A), and for 72h at 37℃ in anaerobic conditions for *S. mutans* and *P. gingivalis* (group B). After incubation the inhibition zones around A-PRF membranes, discs with LP and liquid drops of LP were measured. Tests were done according to Scheme 1.

**Scheme 1.**
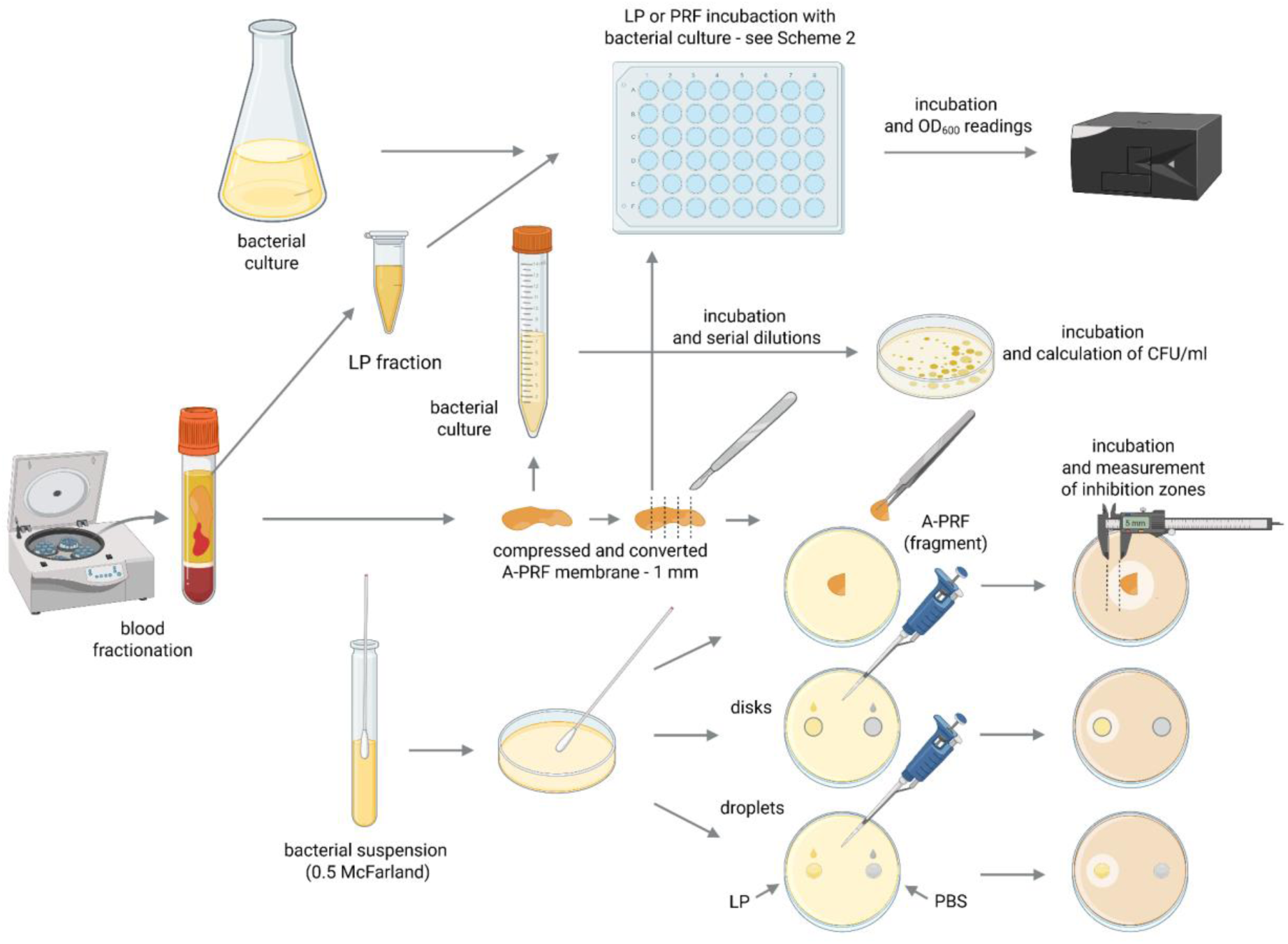
Schematic diagram of the conducted experiments. Created in BioRender (Z, M. (2025) https://BioRender.com/n5zati5). Fractions acquired from blood centrifugation (LP and A-PRF) were used for antibacterial testing performed in liquid (titration plates, OD_600_ spectrophotometer readings) or on solid medium (calculation of CFU/ml and inhibition zones measurements); **LP** - liquid fraction of plasma remaining after blood centrifugation; **A-PRF** - advanced platelet-rich fibrin; **PBS** - phosphate buffered saline.

To perform the test using liquid medium, titration plates (48-well, flat-bottom; Avantor) were prepared following Scheme 2. Briefly, 300 µl of two-fold concentrated Mueller-Hinton Broth (Oxoid) or eTSB culture medium was added to each well. Then, wells were supplemented with either 300 μl of bacterial suspension during logarithmic growth phase (OD_600_ = 0.8 or 0.4) or 300 μl PBS as a control. One fragment of A-PRF membrane (one whole A-PRF membrane was cut into a maximum of 5 smaller pieces of equal size) or 100 µl of LP was added to selected rows of wells with bacterial culture. One row was left as a positive control for bacterial growth. Additionally, sterility control was applied for A-PRF and LP fractions where no bacterial inoculum was added to selected wells (Scheme 2).

**Scheme 2.**
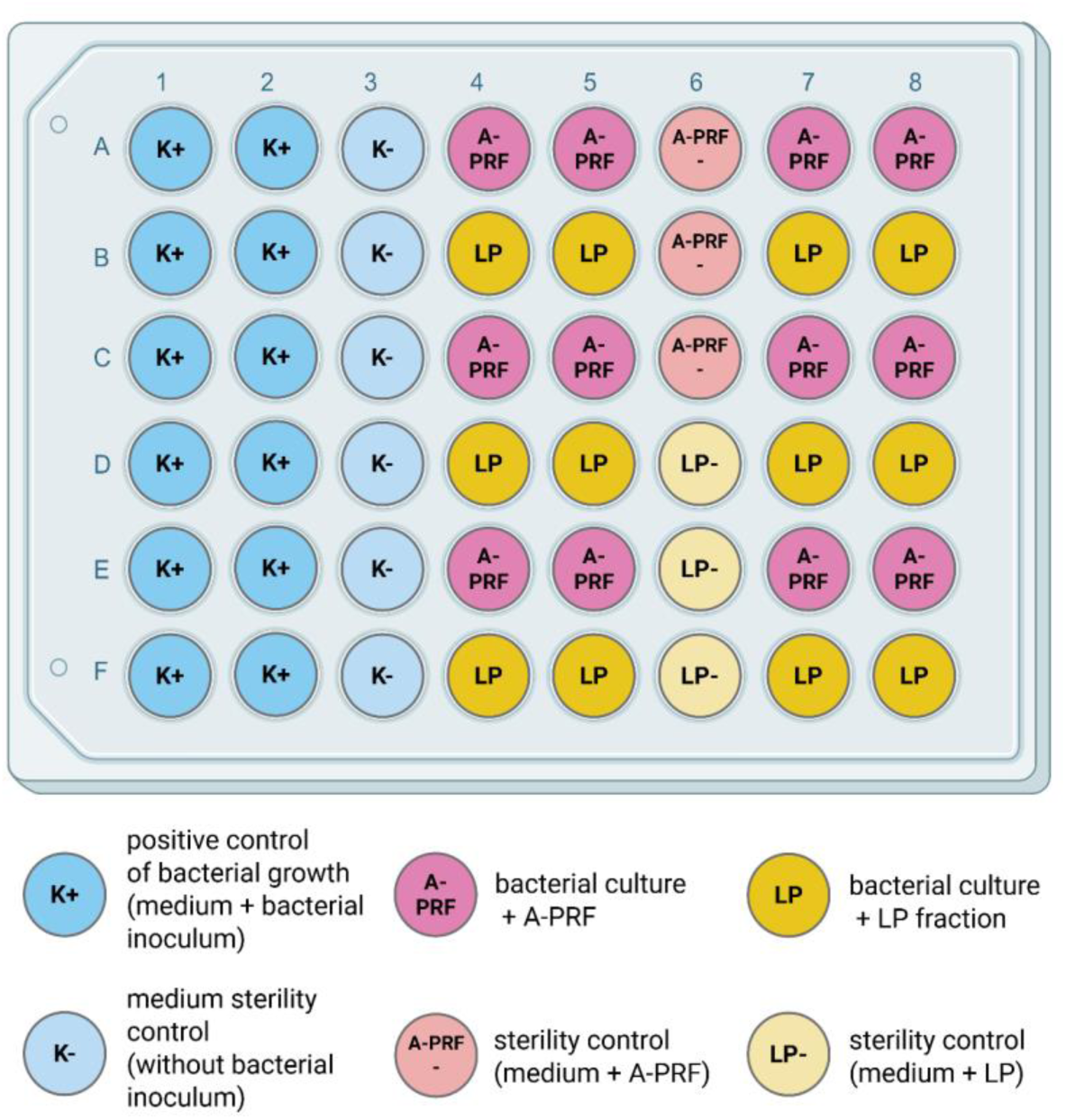
Schematic view of the experiment setup on the titration plate. Created in BioRender (Z, M. (2025) https://BioRender.com/0j45vwx). Antibacterial activity of LP or A-PRF was performed in a liquid medium and assessed by OD600 spectrophotometer readings. One plate was used either for 3 or 1 strain from group A or B, respectively; **LP** - liquid fraction of plasma remaining after blood centrifugation; **A-PRF** - advanced platelet-rich fibrin

Prepared plates were incubated at 37 ℃ in aerobic conditions for *E. coli, B. subtilis, M. luteus, S. lentus, E. casseliflavus* (group A), and in anaerobic conditions for *S. mutans* and *P. gingivalis* (group B). Growth rates were determined by OD_600_ measurements on a spectrophotometer (SpectraMax® iD3 Multi-Mode Microplate Reader, Molecular Devices) every 60 minutes for 7 hours and then after 24 hours (group A) and every 24 hours for 72 hours (group B). Due to the presence of a solid A-PRF matrix in the wells, spectrophotometric measurement of the entire suspension resulted in artificially elevated OD_600_ values. Therefore, for samples containing A-PRF, a portion of the bacterial suspension was carefully removed and measured separately using a spectrophotometer to obtain accurate OD_600_ readings (Berthold Detection Systems GmbH).

For *E. coli*, *S. mutans* and *P. gingivalis* the experiment was repeated using the whole A-PRF membranes (from two patients in each case) in a larger volume (3ml) of MH or eTSB culture medium, respectively. A positive control for the growth of the tested bacterial strains was also prepared (culture without A-PRF). 0.3 ml of an overnight culture of the tested strain was added to each tube. Incubation, in the specified conditions described above, was carried out in Falcon tubes (15 ml) for 24 hours and then a series of dilutions of 10^-1^ – 10^-7^ were prepared and plated (100 µl) on a dedicated solid medium (as described in p. 2.6) in duplicate. Plates were then incubated for 24h - 72h at 37 ℃ in aerobic conditions for *E. coli* and in anaerobic conditions for *S. mutans* and *P. gingivalis.* After incubation, colony forming units (CFU/ml) were determined.

## 4. Conclusions

The study revealed no significant differences in the composition of the oral microbiome between male and female participants assigned to the experimental group; however other factors not included into participants’ survey may comprise differences. Regardless of its form, PRF demonstrated antibacterial activity, with the strongest effect observed against the Gram-negative bacteria *E. coli*, and among Gram-positive species, the highest inhibition was noted for *S. lentus*. In both cases, the antibacterial effect was evident up to 7 h of incubation. For *P. gingivalis*, such an effect was observed only at low initial OD_600_ up to 24 h of incubation. No inhibitory effect of A-PRF membrane was observed against *S. mutans*. Notably, after 24 hours of incubation, complete degradation of the A-PRF membrane occurred in cultures of *P. gingivalis* and *S. mutans*, which was not observed for other tested bacterial strains, which confirms our earlier hypothesis. The LP fraction of PRF exhibited no antibacterial activity against either *P. gingivalis* or *S. mutans*.

These findings indicate that A-PRF membrane therapy should be applied only in the absence of oral infection. In the presence of signs of periodontitis, appropriate antimicrobial treatment should first be administered to eliminate the risk of *P. gingivalis* proliferation, which may compromise the effectiveness and safety of PRF-based therapies.

## Data availability statement

Data sets created during the study covering bacterial and fungal community structure data (raw data from V3-V4 variable regions of 16S rRNA gene and ITS sequencing) have been deposited in publicly available repository under the doi number: https://doi.org/10.58132/J9YP72.

## Author contribution

WP and MP: conceptualization and writing-review and editing, project administration. WP: supervision; DD, DK, RO, MM-G and MP: investigation; MP, MZ and RO: formal analysis and visualization; WP, MZ, AŁ and MP: writing – original draft preparation. All authors read and agreed to the published version of the manuscript.

